# VASP regulates the polar organization of adhesion-associated actin filaments

**DOI:** 10.1101/2025.10.08.681099

**Authors:** Wen-Lu Chung, Rajaa Boujemaa-Paterski, Sabina Winograd-Katz, Matthias Eibauer, Benjamin Geiger, Ohad Medalia

## Abstract

Focal adhesions (FAs) are dynamic macromolecular assemblies that anchor the actin cytoskeleton to the extracellular matrix via integrin receptors, thereby regulating cell morphology and migration. Although FA maturation and organization have been extensively studied, it remains unclear how regulatory proteins influence the 3D architecture of FAs. Here, we show that loss of the vasodilator-stimulated phosphoprotein (VASP) impairs adhesion dynamics. We employed CRISPR/Cas9-mediated knockout of VASP and/or the mechanosensitive adaptor protein zyxin to investigate their respective roles in actin–adhesion coupling. Loss of VASP and zyxin correlates with altered FA morphology and impaired dynamics. Using cryo-electron tomography (cryo-ET), we resolved the polarity of individual actin filaments associated with FAs and identified a contractility-related actin layer enriched with tropomyosin. VASP and zyxin are required for the assembly of dense and aligned actin bundles with uniform polarity, oriented with their barbed ends towards the cell edge. In contrast, the tropomyosin-decorated dorsal actin layer remains unaffected by these perturbations. Our findings reveal distinct, layered architectures within FAs and underscore the cooperative role of VASP and zyxin in stabilizing the organization of actin filaments at functional adhesion sites.

## Introduction

Integrin-based focal adhesions (FAs) are specialized, mechanosensitive cellular organelles that physically link the actin cytoskeleton to the extracellular matrix (ECM). This transmembrane connection enables the generation of sustained mechanical forces, which, in turn, modulate the dynamic assembly and disassembly of both the adhesions and the associated actin filaments (Case and Waterman, 2015; Jansen et al., 2017; Kanchanawong and Calderwood, 2023; Medalia and Geiger, 2010). The tight interplay between key FA-associated proteins, such as talin, vinculin, paxillin, and zyxin was extensively investigated in recent years, focusing primarily on the initiation of FA formation, induced by the mechanosensitive interactions between integrin, talin and vinculin, e.g. (Aretz et al., 2023; Franz et al., 2023; Tapia-Rojo et al., 2023). Nonetheless, the molecular mechanism underlying the tight coordination between actin polymerization at the cells’ leading edge and the formation, maturation and dynamics of adhesions remains an unresolved challenge. Notably, three protein families, enabled/vasodilator-stimulated phosphoproteins (Ena/VASP), zyxin, and tropomyosin, play complementary and interconnected roles in this process. These proteins regulate the structure, function, and mechanical responsiveness of both the actin cytoskeleton and the associated adhesion complexes (Faix and Rottner, 2022; Gunning et al., 2015).

FAs are multi-protein assemblies that form in response to external mechanical cues and internal contractile forces generated by the actomyosin cytoskeleton (Horton et al., 2016; Parsons et al., 2010; Schiller et al., 2011; Winograd-Katz et al., 2014; Zaidel-Bar et al., 2007). Mature FAs are vertically stratified into three distinct layers: the integrin signaling layer (Medalia and Geiger, 2010; Patla et al., 2010), the force transducing layer, and the actin-regulatory layer. The latter is physically linked to actomyosin stress fibers and contains Ena/VASP proteins, zyxin, tropomyosin (Cagigas et al., 2022; Case and Waterman, 2015; Crawford et al., 1992; Gateva et al., 2017; Kanchanawong et al., 2010; Legerstee and Houtsmuller, 2021; Rottner et al., 2001; Uemura et al., 2011; Valencia et al., 2021). Studies have further shown that mechanical stimulation induces a retrograde flow of zyxin and VASP from FAs to the attached actin stress fibers, which may function as a mechanical clutch for cell migration (Guo and Wang, 2007; Yoshigi et al., 2005). However, the extent to which these proteins modulate the architecture of adhesion-associated actin networks remains elusive.

Ena/VASP family proteins, comprising Evl, Mena and VASP, are actin elongation factors that promote the formation of long, unbranched filaments by antagonizing capping proteins and facilitating barbed-end polymerization (Breitsprecher et al., 2011; Hansen and Mullins, 2010; Rottner et al., 2001). These proteins localize to FAs and to the tips of lamellipodia, where they sustain actin filament elongation and retrograde actin flow (Bieling and Rottner, 2023; Hansen and Mullins, 2015; Kage et al., 2022; Lavelin et al., 2013). Among the three, Evl has been identified as the major regulator of FA formation, indirectly controlling cell spreading, steering, and actomyosin traction force generation. In contrast, Mena and VASP are thought to have a more limited role in these processes (Damiano-Guercio et al., 2020; Puleo et al., 2019; Rottner et al., 2001). VASP was shown to effectively regulate the actin meshwork in lamellipodia, its architecture, width and the process of cell motility (Damiano-Guercio et al., 2020). Physiologically, triple knockout (KO) mice lacking all Ena/VASP family members exhibit embryonic lethality at E16.5 stage, accompanied by defects in neurite outgrowth (Kwiatkowski et al., 2007), highlighting the central roles of Ena/VASP proteins in developmental processes and cytoskeletal dynamics.

It should be noted that FA maturation depends on distinct mechanosensitive proteins (Askari et al., 2010; Schiller and Fassler, 2013). One of these is Zyxin, a LIM domain protein that functions as an adaptor at FAs and stress fibers. Its three C-terminal LIM domains enable zyxin to cluster at FAs and stress fibers in response to mechanical stress (Hirata et al., 2008). Zyxin N-terminus contains α-actinin binding sites and proline-rich domains, which recruit Ena/VASP proteins and are crucial for maintaining stress fiber integrity (Drees et al., 2000; Hansen and Beckerle, 2006; Wang et al., 2019). Similar to vinculin, a mechanosensing FA protein with proline-rich domains, zyxin anchors VASP to FAs, directing the actin polymerization machinery at these sites (Drees et al., 2000; Zaidel-Bar et al., 2003). Loss or mis-localization of zyxin disrupts Ena/VASP positioning at FAs and impairs stress fiber remodeling (Ghosh et al., 2015; Hoffman et al., 2006; Li and Burridge, 2019; Smith et al., 2010), resulting in disorganized actin networks and reduced protrusion stability. Physiologically, zyxin knockout mice show no obvious phenotype (Al-Hasani et al., 2022; Hoffman et al., 2003), suggesting possible functional compensation or redundancy (Hoffman et al., 2006).

Taken together, these studies suggest that Ena/VASP, zyxin, and tropomyosins concertedly ensure protrusive force generation and adhesion maturation to be temporally and spatially coordinated. Ena/VASP promotes actin elongation at the leading edge and at FAs, supporting their growth and strengthening. Zyxin functions as a tension-sensitive scaffold that spatially organizes and recruits Ena/VASP, and tropomyosins regulate actin filament mechanical stability and support actomyosin contractility (Faix and Rottner, 2022; Hardeman et al., 2020; Oldenburg et al., 2015; Tojkander et al., 2011). However, the precise molecular mechanisms by which these proteins regulate FA-associated actin fibers remain unclear. Here, we show that, unlike zyxin, loss of VASP impairs adhesion dynamics while loss of either protein alters FA distribution and stress fiber organization. By using cryo-electron tomography (cryo-ET), we explore the nano-architecture at the FA-stress fiber interface and elucidate the involvement of VASP in the organization of FA-associated actin, reveal the association of tropomyosin with the dorsal aspect of the FA-bound stress fiber, and resolve actin filament polarity at FAs in the intact wild-type, VASP^-/-^, zyxin^-/-^, and double knockout (DKO) mouse embryonic fibroblasts (MEFs). Loss of VASP leads to altered FA morphology, reduced density of FA-associated actin, and significantly disrupted filament polarity near the plasma membrane, a defect that is more pronounced in zyxin^-/-^ and DKO cells. Tropomyosin decoration of actin filaments appears to be independent of VASP and zyxin. Thus, while VASP and zyxin may be mechanistically linked, they display distinct roles in coupling adhesions to the actin cytoskeleton and regulating their dynamics.

## Results

### Loss of VASP alters adhesion dynamics

We previously reported that actomyosin contractile forces regulate the composition of FAs, and that their mechanosensitivity selectively modulates the on- and off-rates of individual FA components. Particularly, under low or no-force conditions, newly formed peripheral FAs lack VASP and zyxin, and both proteins are among the fastest to dissociate during adhesion disassembly (Lavelin et al., 2013). To further investigate the role of these proteins in FAs, we analyzed adhesion dynamics in MEFs expressing vinculin-Venus (Grashoff et al., 2010), and sequentially knocked out VASP and zyxin using the CRISPR/Cas9 technology (Fig. 1a; Fig. S1). Cells were plated on fibronectin-coated glass surfaces, and time-lapse series were acquired at an interval of one-minute with fluorescence microscopy. Adhesion structures were segmented as particles of different sizes, and comparison between the initial and the following frames was used to define distinct regions within the adhesion structure. This approach enabled us to quantify persistent, newly formed, and lost FA areas, using the fluorescence signal. WT and mutant cell lines exhibited significant, albeit subtle, differences in adhesion size, consistent with the wide distribution of adhesion areas observed (Fig. 1b, S2a-d). Next, we analyzed adhesion dynamics based solely on pixel-fluorescence information. We compared pixel fluorescence intensity between the first frame and subsequent frames, representing later time points (Fig. 1c, d), or used these frames to perform autocorrelation analysis (Fig. S3). The decline in counts of individual pixels was nonlinear and continued over time (Fig. 1c), likely due to *de novo* recruitment of proteins to the adhesion site that subsequently persisted. Interestingly, cells lacking VASP displayed the highest proportion and slowest decline of persistent signals (Fig. 1c), although the rates of newly formed and lost signals were comparable to those in WT cells (Fig. 1d). Conversely, DKO cells exhibited the lowest proportion of persistent signals (Fig. 1c), correlating with the lowest rate of newly formed adhesions (Fig. 1d). In contrast, the loss of zyxin did not significantly affect adhesion dynamics compared to WT (Fig. 1c, d), in line with previous work (Hoffman et al., 2006). Thus, the loss of VASP slows vinculin turnover at adhesion sites, whereas the combined loss of VASP and zyxin has the opposite effect, accelerating vinculin exchange and reducing adhesion stability.

**Figure 1.**
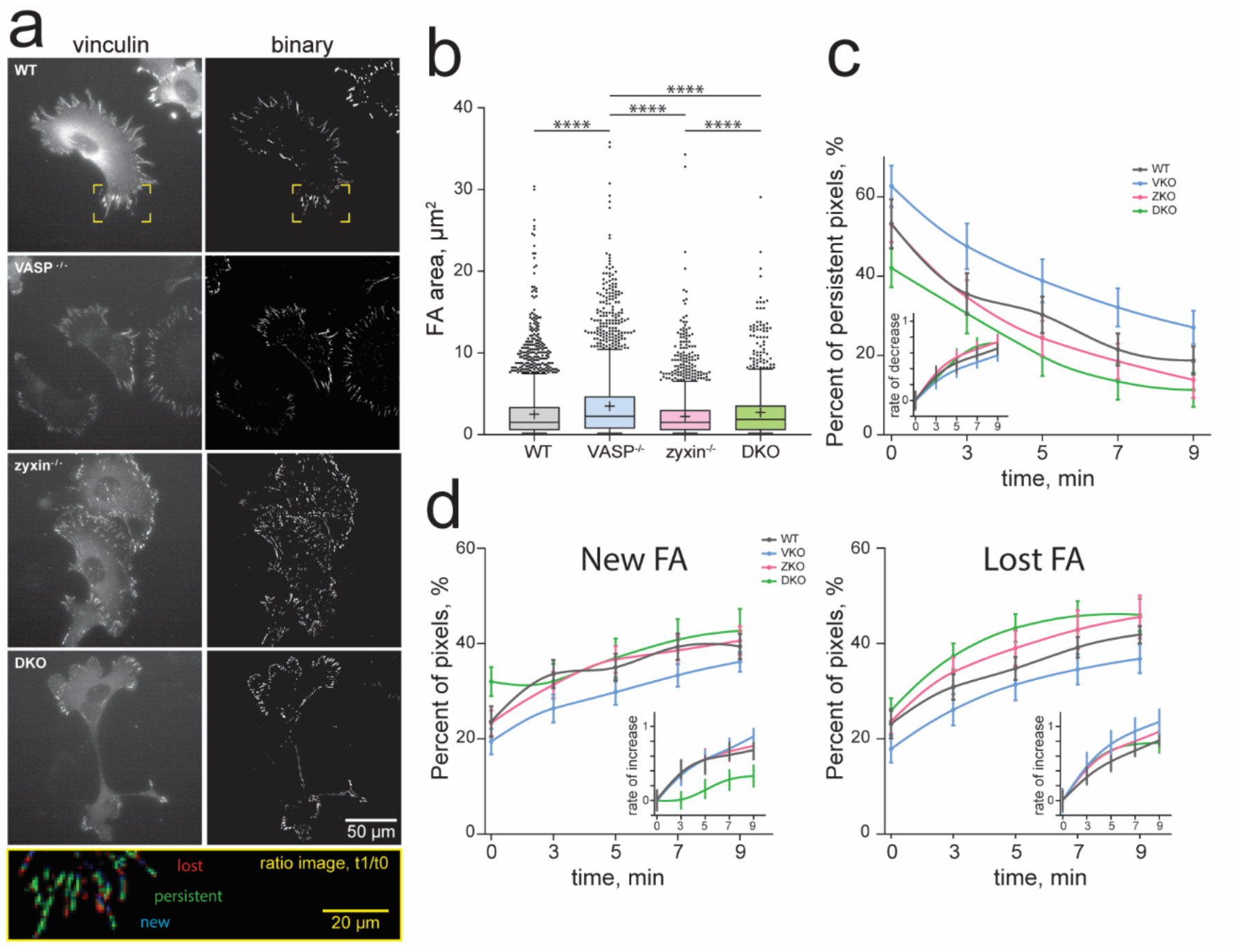
Loss of VASP alters adhesion dynamics in vivo. **a**, Representative images of cells spread fibronectin-coated surfaces, and corresponding binary image obtained after adhesion fluorescent pixel classification using Ilastic software (see Methods). As illustrated for the yellow region highlighted in the WT image, for each cell line, temporal ratio images were generated by dividing the intensity of each pixel by its intensity at the initial time point. In these images, adhesion pixels appear green when persistent, blue when newly formed, and red when lost. **b,** Quantification of the area of segmented FAs observed in each cell line. Statistical comparisons using the Holm-Šídák test, and a one-way analysis of variance (ANOVA) showed significant variations among the measured FA parameters, with p values **** = <0.0001. The number of analyzed cells was 6 for WT, 6 for VASP^-/-^, 6 for zyxin^-/-^, 7 for DKO. Measurements were taken from at least two distinct samples per cell line. **c-d,** For each cell line, the quantification of persistent, new and lost pixels are represented over time. Inserts show the rate of decrease (**c**) or increase (**d**) from the start. Data points represent mean values, bars, standard deviation, and smoothing spline fits (number of knots = 4) were plotted using GraphPad prism. (see related Fig. S1-3)

### Loss of VASP and zyxin alter distribution of FAs and stress fiber organization

To further characterize the impact of VASP and zyxin on FA formation, we utilized circular micropatterns on glass surfaces to spatially confine cell shape, thereby controlling the distribution and morphology of FAs (Tee et al., 2015). The geometric confinement directs adhesions to the periphery of the adhesive area, enabling a quantitative and more accurate analysis of FA morphology and distribution in the WT, VASP^-/-^, zyxin^-/-^, and DKO MEFs (Fig. 2a, upper row). Additionally, the circular patterns impact the organization of the actin cytoskeleton, prominently highlighting dorsal stress fibers that originate at the adhesion sites and connect to radial contractile fibers, which flow centripetally (Fig. 2a, upper row) (Tee et al., 2015). Due to the high actin network density, quantifying individual stress fibers was challenging (Fig. 2a, upper row, Fig. S4). However, the spatial organization of segmented FAs could be represented using averaged images of more than 100 patterned cells (Fig. 2a, lower row, and Fig. 2b). Loss of VASP led to reduced lamellipodial and ruffle extensions, consistent with previous findings (Damiano-Guercio et al., 2020), while zyxin knockout resulted in a marked reduction in FA size and vinculin density without affecting the overall number of adhesions, suggesting impaired FA maturation (Fig. 2c-f). Knockout of VASP, zyxin, or both impacts negatively the density and number of dorsal and radial fibers, in line with the significantly smaller FAs observed (Fig 2f). Stress fibers appeared thinner and shorter, while membrane protrusions remained prominent (Fig. S4), indicative of diminished contractility and disrupted actin remodeling, as previously reported (Smith et al., 2010). Notably, DKO cells exhibited additive defects compared to the individual knockouts, underscoring a cooperative role for VASP and zyxin in regulating FA-associated cytoskeletal architecture.

**Figure 2.**
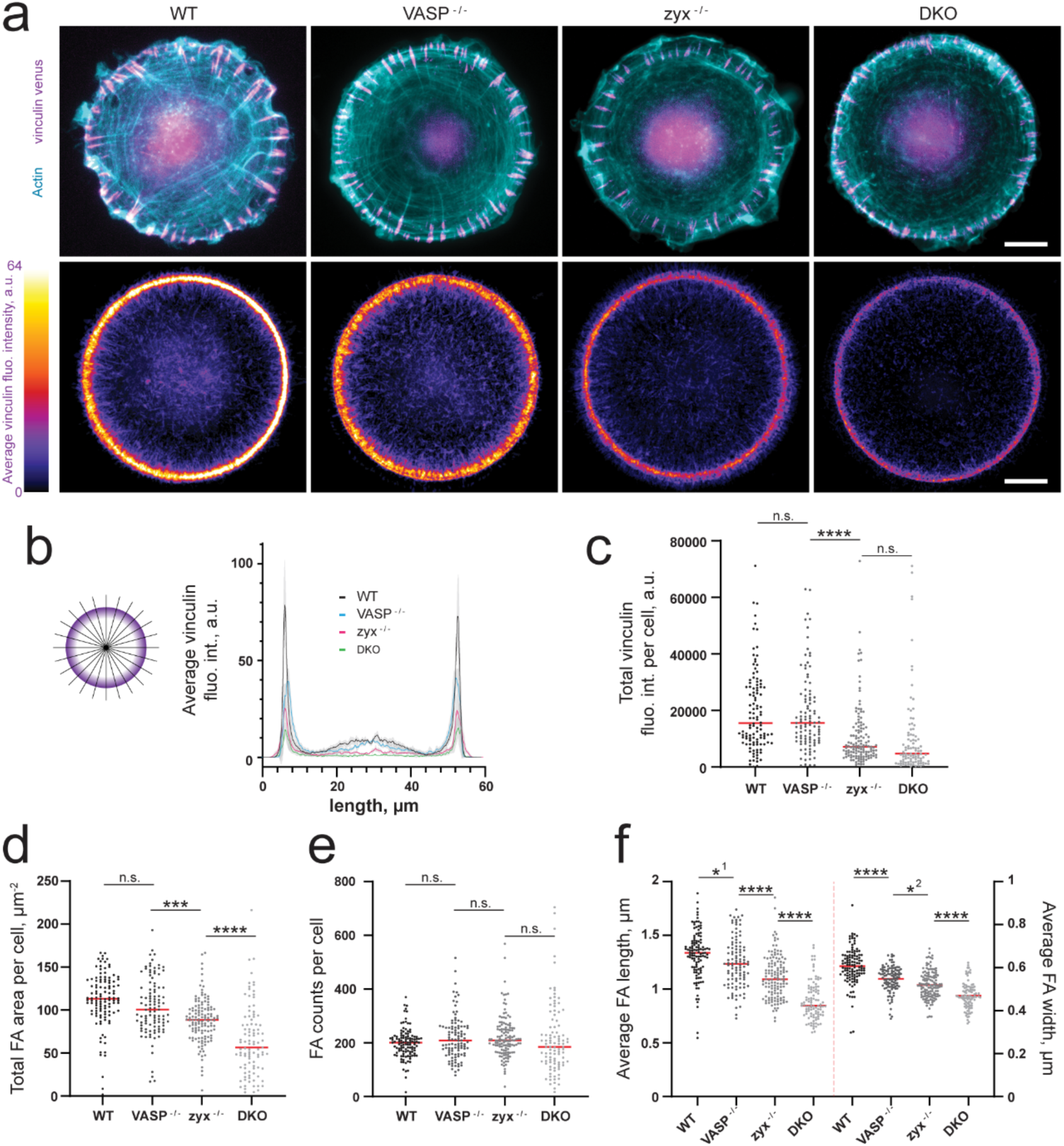
Loss of VASP and zyxin affects FA organization of cells on patterned surfaces. **a**, Representative images of cells spread on circular patterned surfaces (∼1810 µm^2^, top row), average distribution of vinculin fluorescence intensity obtained by superimposing of patterned cells, 112 for WT, 104 for VASP^-/-^, 129 for zyxin^-/-^, 96 for DKO (bottom row). Measurements were taken from at least two distinct samples per cell line. Vinculin: magenta; actin: cyan. Scale bar is 10 µm **b,** Vinculin fluorescence intensity was measured along radial lines (as shown the diagram) drawn on the average images in **a**. Bars in grey represent the standard deviation. **c-f,** Quantification of the total cellular vinculin fluorescence intensity (**c**), the total cellular FA area (**d**), cellular FA number (**e**), and FA morphology (**f**) for each cell line and using the average images in **a**. Red lines are the mean values. Statistical comparisons using the Holm-Šídák test, and a one-way analysis of variance (ANOVA) showed significant variations among the measured FA parameters, with p values for **c-f**: **** = <0.0001 / *** = / *1 = 0.0287 / *2 = 0.0103 / n.s. = 0.9971; 0.9768; 0.3365; 0.4902; >0.9999; 0.9897. (see related Fig. S4)

### Tropomyosin-decorated actin filaments at FAs

To gain higher-resolution insight into the polarity of actin filaments interfacing FA sites, we applied cryo-ET to FAs (Elad et al., 2013; Patla et al., 2010), using actin polarity determination approach. This method previously demonstrated that actin bundles predominantly orient their barbed ends toward the cell margin and exhibit more uniform filament polarity than the membrane-distal FA region, located towards the cell body (Martins et al., 2021).

Cells were seeded on fibronectin-coated gold EM grids for 3h to allow FA formation prior to live fluorescence imaging and vitrification. At this time point, FAs had formed, facilitating cryo-ET data acquisition and analysis. The peripheral regions of MEFs are sufficiently thin to allow cryo-ET imaging of intact cells. Correlative light and electron microscopy (CLEM) guided the cryo-ET acquisition, to ensure relevant datasets (Fig. 3a). Tomograms were then acquired from membrane-proximal regions of each FA (Fig. 3a, white frames), from ≥8 cells per cell line (45 for WT, 8 for VASP^-/-^, 29 for zyxin^-/-^, and 13 for DKO). Characteristic bundled actin filaments, alongside single filaments and macromolecular complexes, were visible in raw *x-y* slices of each tomogram (Fig. 3a). The high quality of these datasets enabled automated 3D filament annotation using CrYOLO (Wagner et al., 2019). Next, the Actin Polarity Toolbox (APT) (Chung et al., 2022) was employed to extract subtomograms of actin filaments (Fig. S5a), yielding a reconstructed actin filament structure at 12 Å resolution (Fig. 3b-d, S6a-c). At this resolution the polarity of each filament can be reliably determined. Interestingly, a subset of tropomyosin-decorated actin filaments was identified during 2D classification (Fig. S6a, red boxes). These filaments were iteratively re-extracted and added to the particle pool, ultimately constituting ∼6% of all analyzed actin filaments. (Fig. S6b). The final reconstructed tropomyosin-actin filament was resolved at 13 Å, consistent with previously published structure (Wang et al., 2021) (Fig. 3e-g, Fig. S6d). Thus, this cryo-ET analysis enabled a detailed actin filament polarity mapping within FAs and resolved tropomyosin-decorated actin filaments at adhesion sites. Tropomyosins have previously been shown to be present in FAs (Kumari et al., 2024).

**Figure 3.**
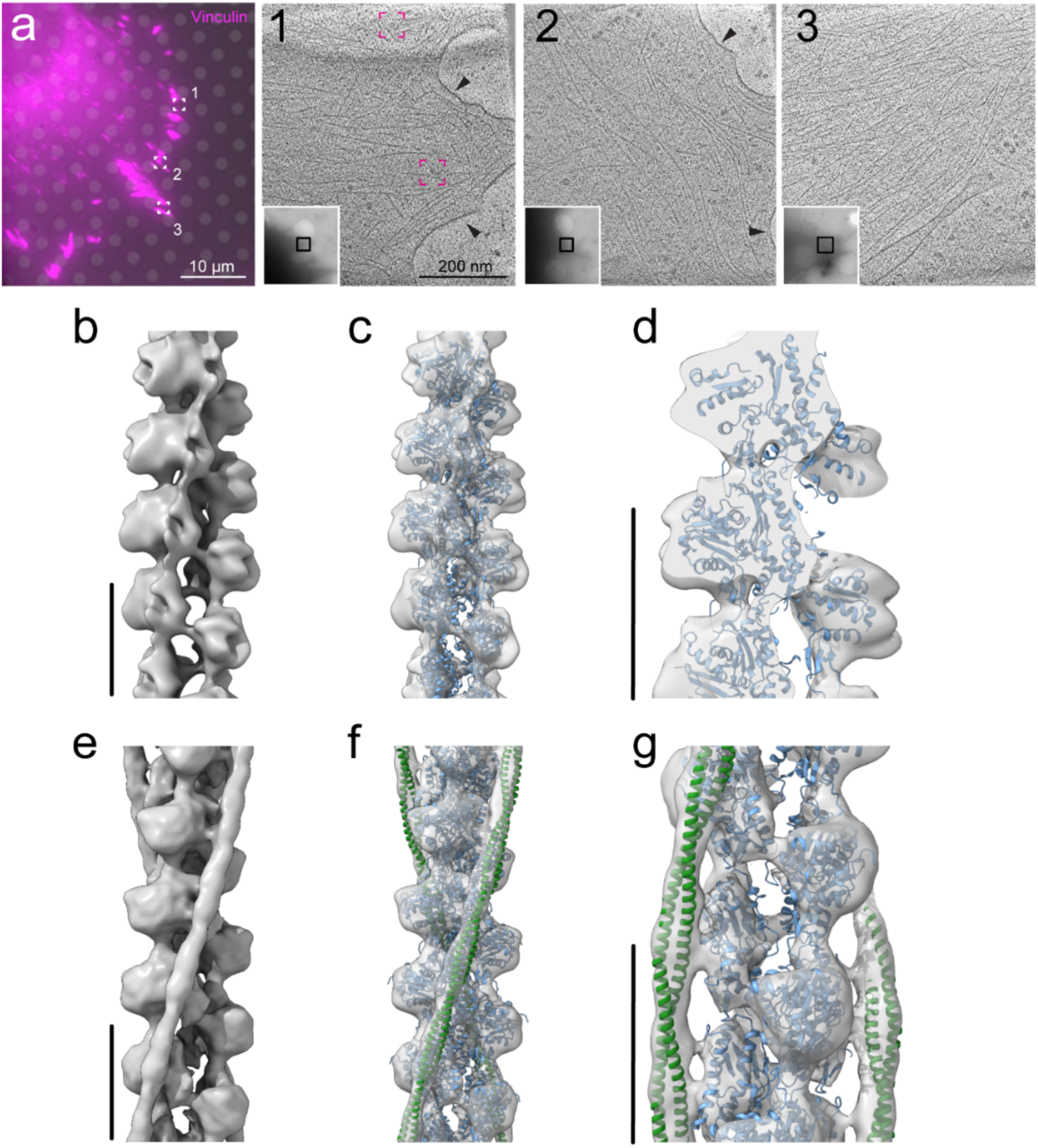
Three-dimensional reconstruction of actin filament and Tropomyosin-actin filament from focal adhesion-associated actin networks. **a** A representative image of a WT cell adhered to an EM grid, expressing fluorescently tagged vinculin-Venus (magenta). Areas selected for cryo-ET analysis are indicated by boxes 1–3. **1–3,** CLEM: Representative x–y slices from each tomogram acquired within the regions boxed in black in the low-magnification EM image (lower right), corresponding to the boxed areas in panel a. Magenta boxes encircle single or multiple adhesion-associated macromolecular complexes, and arrowheads indicate the double leaflet of the plasma membrane **b,e** Structure of actin filament and tropomyosin-actin filament resolved 11.4 Å and 13 Å, respectively. **c,f** Actin filament’s barbed end and pointed end were identified by docking the canonical F-actin structure (EMD-15106) and the canonical Tropomyosin-F-actin structure (EMD-12289), respectively. **d,g** Enlarged views of panels c and f, highlighting the high-resolution reconstruction of actin and tropomyosin-decorated actin filaments, respectively. Scale bars are 8 nm. (see related Fig. S5-6)

### Actin polarity at FAs of WT and cells deficient in VASP and zyxin

Back-mapping of the actin filaments was conducted to reveal their polarity distribution and spatial organization, as previously described (Martins et al., 2021). We applied this approach to FAs in WT and mutated cells (Fig. 4a and S7). Actin filaments with barbed ends oriented toward the cell plasma membrane (Fig. 4a) and aligned along the FA axis (Fig. 4b) were classified as ‘forward’ filaments (blue colored actin filaments in Fig. 4 and Fig. S7), while those oriented toward the cell body were designated as ‘backward’ filaments (red colored actin filaments in Fig. 4 and Fig. S7). Transverse filaments, which were not aligned with the FA axis, were occasionally observed and colored in brown. Cell substrates were manually segmented in each tomogram and aligned to the z = 0 plane (Fig.4c, d and S5b, magenta arrowhead).

**Figure 4.**
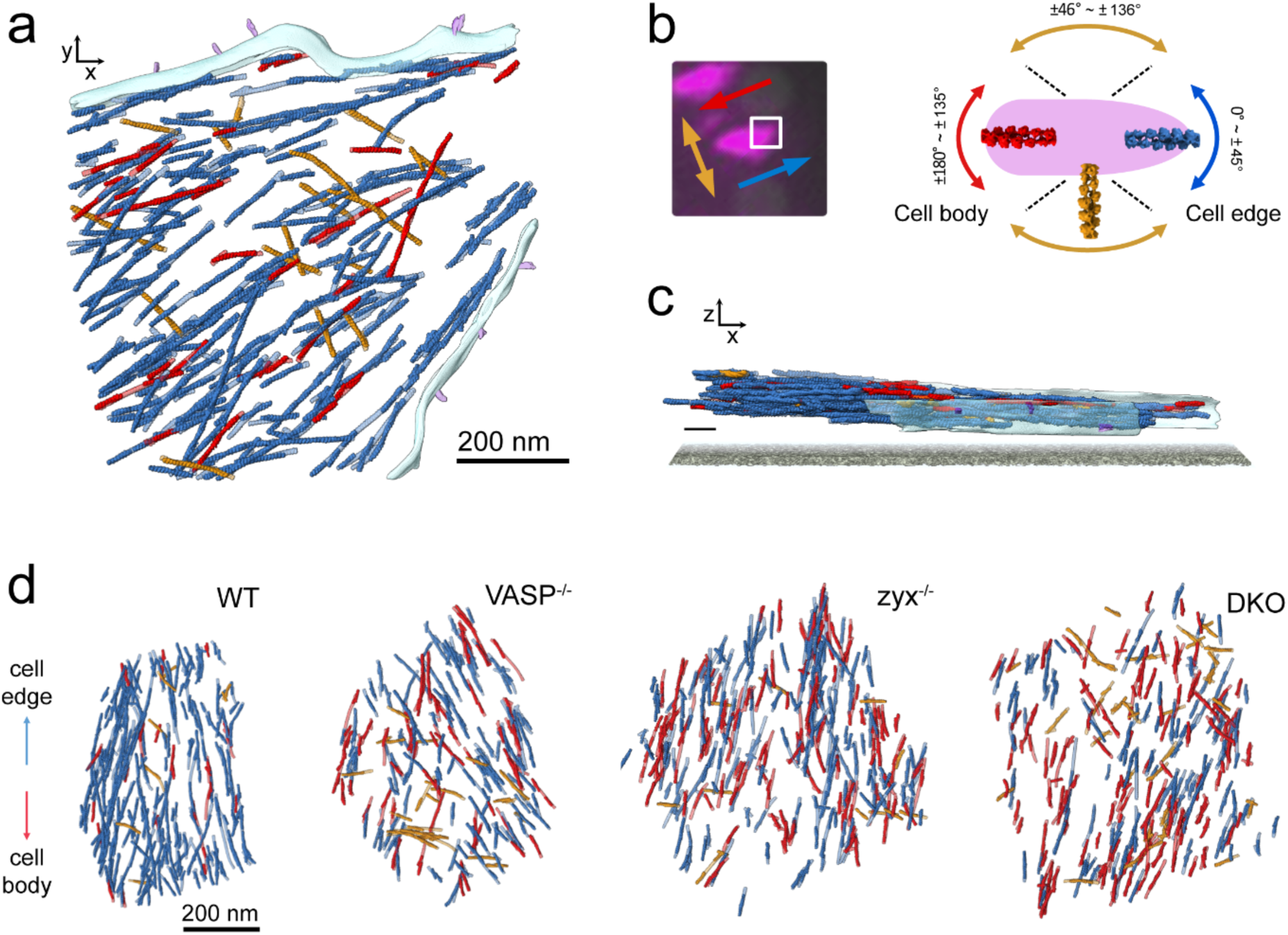
Elucidation of the filament network and actin polarity at FA. **a** An x-y surface-rendered view of a tomogram taken from a WT cell. Actin filaments are color-coded according to their orientation relative to the FA main axis and the plasma membrane (color scheme shown in **b**). The plasma membrane was manually segmented and is shown in light blue. **b** A fluorescence image showing the adhesion tip where cryo-ET was applied (white box). Actin filament orientation is indicated by arrows and color-coded as depicted in the schematic: forward filaments (blue) have their barbed ends oriented toward the cell periphery along the FA axis; backward filaments (red) are oriented in the opposite direction; transverse filaments (brown) are not aligned with the FA axis. **c** A z-x surface-rendered side view of panel a. in which the cell substrate was manually segmented in the original tomogram and aligned to the z = 0 plane. Scale bar is 50 nm. **d** Representative x-y surface-rendered views for WT and mutant cells. (see related Fig. S4b and S7).

To characterize the role of VASP and zyxin in the organization of adhesion-associated actin filaments, we analyzed individual and combined KO cells using cryo-ET (Fig. S7) and determined the polarity of the cytoskeleton network (Fig. 4d). While the positions and angles of the actin filaments relative to the substrate were similar across all cell lines (Fig. 5a, b), the polarity and density of actin filaments within FAs appeared substantially altered by the lack of VASP and zyxin (Fig. 5c, d). In WT cells, the densely packed filaments exhibited a high degree of uniform polarity, with 71% oriented forward. In contrast, a decrease in forward filaments was detected in VASP^-/-^, zyxin^-/-^, and DKO cells (68%, 65%, 63%, respectively). This was accompanied by an increase in backward filaments, with average enrichments of 10%, 20%, and 26% over the WT cells in VASP^-/-^, zyxin^-/-^, and DKO cells, respectively (Fig. 5c). Interestingly, the density of actin filaments at FAs was not reduced in VASP^-/-^ cells, remaining at ∼4000 filaments per μm^3^. However, it was substantially reduced in zyxin^-/-^ and DKO cells to ∼3000 and ∼2500 filaments per μm^3^, respectively (Fig. 5d). Remarkably, this reduction in filament density in zyxin^-/-^ and DKO cells, representing a decrease of 27% and 32%, respectively, is consistent with the observed reduction in vinculin signal (Fig. 2). Overall, zyxin^-/-^ cells displayed more substantial phenotypic changes than VASP^-/-^ cells, while DKO cells exhibited additive effects. However, the smaller differences in actin polarity between WT and VASP^-/-^ that were detected, are likely due to compensatory activity from other Ena/VASP family members (Damiano-Guercio et al., 2020; Puleo et al., 2019).

**Figure 5.**
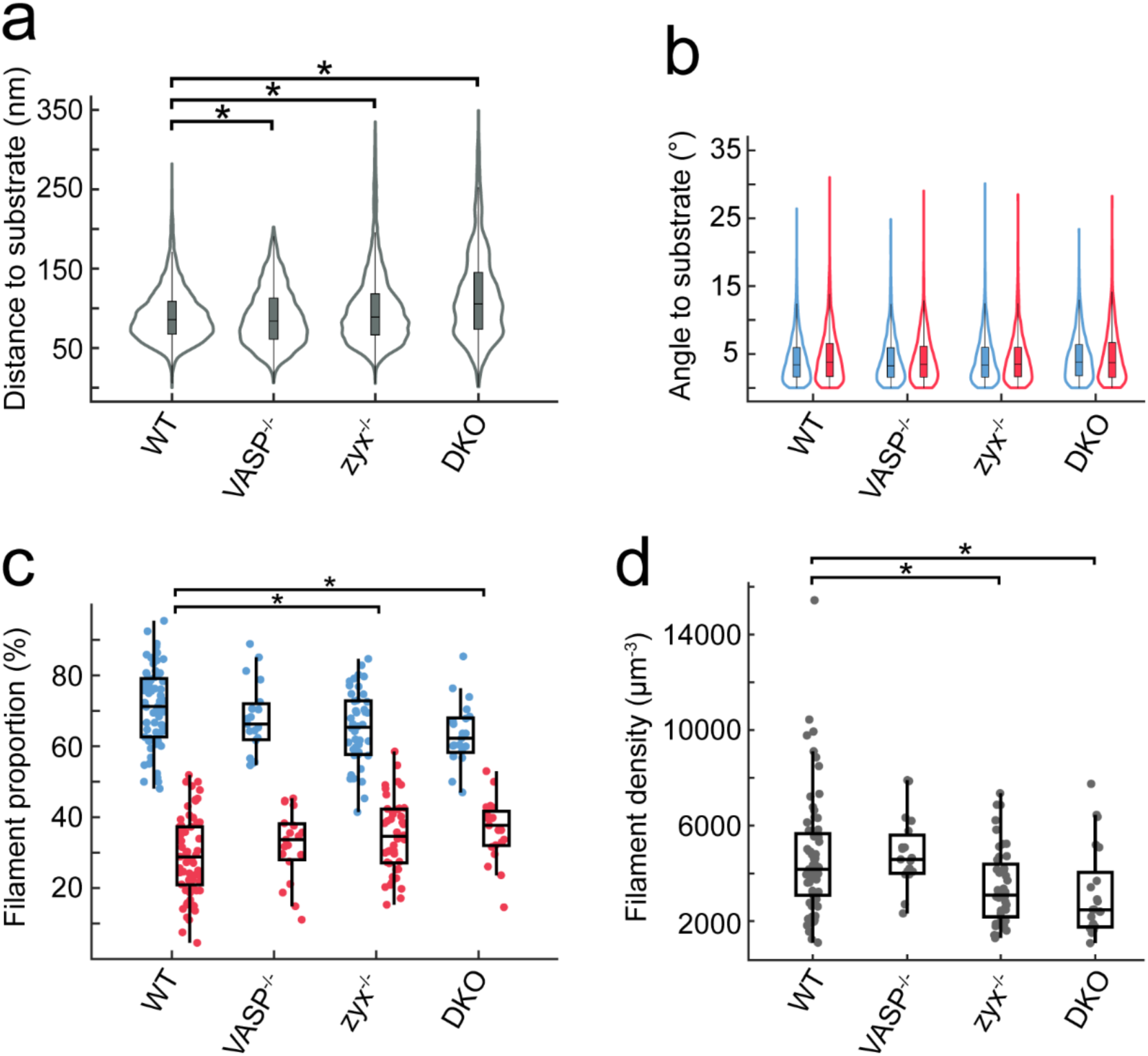
Knockout of VASP and zyxin affects actin polarity establishment and filament density at FAs. **a** Violin plot representing the distribution of distances from the center of mass of actin filaments to the substrate. **b** Violin plot representing the distribution of angles to substrate measured for forward (blue) and backward (red) actin filaments. The box within each violin plot indicates the interquartile range (25^th^ to 75^th^ percentiles), with the horizontal line representing the median (50th percentile), and the whiskers extending to the 10^th^ and 90^th^ percentiles (**a** and **b**). **c** Proportion of forward (blue) and backward (red) filaments. **d** Actin filament density per cube micrometers. Box plots show the data quartiles, with whiskers representing values within 1.5 times the interquartile range (IQR) (**c** and **d**). Parameters were measured in all cells, using 57 tomograms for WT, 19 for VASP^-/-^, 44 for zyxin^-/-^, 21 for DKO. Measurements were taken from at least two distinct samples per cell line. Statistical comparisons using the Tukey-Kramer test and a one-way analysis of variance (ANOVA) showed significant variations among the measured parameters, with p values \**P* < 0.05. (see related Fig.S8)

### Tropomyosin is associated exclusively with thy dorsal aspect of adhesion-associated stress fibers

Tropomyosins, coiled-coil actin-binding proteins, further fine-tunes this dynamic system. It decorates actin filaments, specifying filament identity by stabilizing specific actin architectures while excluding organizations such as Arp2/3-nucleated branches (Gunning and Hardeman, 2017; Gunning et al., 2015). In lamellipodia, the tropomyosin isoforms Tpm1.8/1.9 are thought to decorate a subset of filaments, often those elongated by formins or Ena/VASP, and contribute to nascent adhesion assembly (Cagigas et al., 2022), while isoform Tpm1.6 contribute to FA maturation and connection to dorsal stress fibers, isoform Tpm3.2 regulate FA turnover (Kumari et al., 2024), and isoform Tpm4 promotes dorsal stress fiber assembly by recruiting myosinII (Tojkander et al., 2011). As such, in FAs, tropomyosin isoforms are key to lamellipodia dynamics, focal adhesion formation and turnover, and traction force generation.

Surprisingly, we found that most tropomyosin-decorated actin filaments were localized to the dorsal aspect of the adhesion-associated actin bundle (Fig. 6a), positioned approximately 20 nm above the center of the actin volume. Tropomyosin signal was predominantly detected at ∼110 nm above the substrate (Fig. 6b, c). The density of tropomyosin-actin filaments (Fig. 6d) was estimated by combining the proportion of tropomyosin-positive filaments (Fig. 6e) with the total actin filament density (Fig. 5d). These filaments were also detected in the KO cell lines, where they remained localized to the upper portion of the actin network (Fig. 6b). Interestingly, although the absolute density of tropomyosin-decorated filaments remained unchanged (Fig. 6d), the relative proportion of decorated filaments appeared elevated in zyxin^-/-^ and DKO cells (Fig. 6e), due to the overall reduction in actin filament density in these mutants (Fig. 5d and S8a). The distribution of tropomyosin among differently oriented filaments mirrored the underlying distribution of actin filament orientations, regardless of genotype or subregion (Fig. S8b). These findings suggest that, at FAs, tropomyosin localization is regulated independently of actin filament polarity and the presence of either VASP or zyxin.

**Figure 6.**
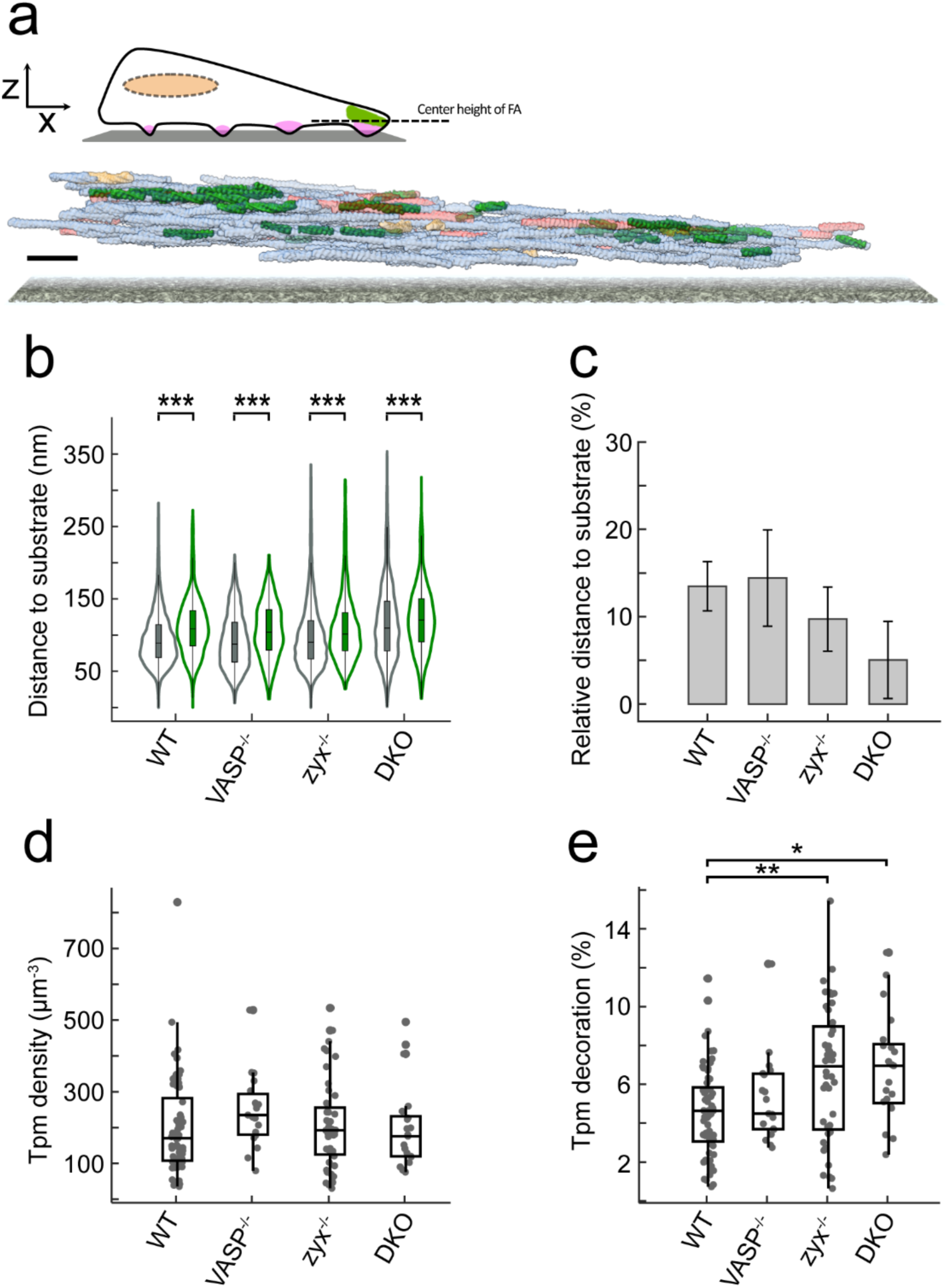
Tropomyosin decorates the dorsal focal adhesion-associated actin. **a** A schematic of an adherent cell depicting the localization of tropomyosin (green) at the dorsal aspect of focal adhesions (magenta), alongside a side view of a rendered view of a tomogram of a FA from a WT cell. Tropomyosin-decorated actin filaments are shown in green, while bare forward and backward filaments are semi-transparently colored in light blue and light red, respectively. The cell substrate was manually segmented in the original tomogram and aligned to the z = 0 plane. Scale bar: 50 nm. **b** Violin plot representing the distribution of distances from the center of mass of tropomysin-actin filaments to the substrate. **c** Relative distance to the substrate of tropomyosin-actin compared bare filaments. **d** Tropomyosin-actin filament density per cube micrometers. **e** Tropomyosin decoration ratio in percentage. The box within each violin plot indicates the interquartile range, with the horizontal line representing the median. Parameters were measured in all cells, using 57 tomograms for WT, 19 for VASP^-/-^, 44 for zyxin^-/-^, 21 for DKO. Measurements were taken from at least two distinct samples per cell line. Statistical comparisons using the Tukey-Kramer test and a one-way analysis of variance (ANOVA) showed significant variations among the measured parameters, with p values: \**P* < 0.05, \*\**P* < 0.01, \*\*\**P* < 0.001. Boxplots show the data quartiles and whiskers of values within 1.5 IQR (**b,c,e**).

## Discussion

Cell adhesions are essential sub-cellular domains that allow cells to sense and respond to mechanical cues from the underlying ECM (Medalia and Geiger, 2010). This responsiveness is mediated largely through the generation and regulation of tensile forces by the contractile actomyosin cytoskeleton. Accessory proteins, including actin cross-linkers like α-actinin and filamin, polymerization regulators like Ena/VASP family members, and tension-dependent zyxin, play crucial roles in maintaining FA-associated stress fibers and fine-tuning of cellular forces (Gateva et al., 2017; Gateva et al., 2014; Lehtimaki et al., 2021; Uemura et al., 2011; Vignaud et al., 2021). Here, using genetic manipulations, combined with fluorescence microscopy, cryo-electron tomography and advanced morphometry, we revealed the impact of VASP on the architecture of focal adhesion-associated actin filaments and uncovered the confined spatial distribution of tropomyosin-decorated actin in FAs at peripheral regions of cells. Advances in cryo-ET have enabled detailed insights into actin directionality in FAs. Despite the abundance of actin binding proteins at FAs, previous *in situ* cryo-ET structural analysis have not resolved the actin binding proteins using conventional image processing approaches. However, due to the relative high occupancy of tropomyosin-decorated actin, these filaments appeared within the structural classes, enabling high-resolution reconstruction and analysis of their 3D structure.

### Integration of filament orientation and lamellipodial flow by VASP at adhesions

VASP and zyxin display similar binding dynamics, which differ from other early adhesion components such as talin, vinculin, and paxillin, mechanosensitive scaffolding proteins (Franz et al., 2023; Lavelin et al., 2013; Legerstee et al., 2019). It is well established that talin, vinculin and paxillin regulate the early recruitment of proteins to nascent adhesions, while many binding partners of the vinculin tail domain, including VASP and zyxin, are only engaged at later FA stages of maturation (Brindle et al., 1996; Drees et al., 2000; Nix et al., 2001; Reinhard et al., 1996; Rottner et al., 2001). Notably, the vinculin tail shares several interaction partners with VASP and zyxin, including each other (Humphries et al., 2007; Zaidel-Bar et al., 2007).

Using knock-out cell lines, our study demonstrated that among Ena/VASP family members (Damiano-Guercio et al., 2020; Faix and Rottner, 2022), VASP plays a role in organizing FA-associated actin. Loss of VASP altered the uniform polarity of the actin network by reducing the proportion of forward-oriented (barbed-end facing) filaments at the cell margin. Although the vertical positioning of the actin network relative to the membrane, and presumably to adhesion complexes, remained unaffected. Moreover, VASP deletion significantly slowed vinculin turnover at FAs. However, in agreement with previous findings, loss of VASP only moderately reduced vinculin recruitment and FAs, in comparison to wild-type cells. FAs appeared slightly narrower and less elongated in the absence of VASP and showed a dramatic reduction in lamellipodial actin (Damiano-Guercio et al., 2020).

A plausible explanation is that VASP regulates retrograde lamellipodial actin flow, thereby contributing to actin polarity at mature adhesions. In this model, VASP promotes efficient lamellipodial actin polymerization, which supports retrograde flow and facilitates the recruitment and/or activation of mechanosensors such as talin, α-actinin, and vinculin to nascent adhesions (Boujemaa-Paterski et al., 2020; Choi et al., 2008; Franz et al., 2023; Kelley et al., 2020). Activated vinculin, in turn, recruits VASP to enrich adhesions with force productive forward-oriented actin filaments (Mogilner and Oster, 1996; Mogilner and Oster, 2003), reinforcing the actin network and enabling clutch engagement with actomyosin stress fibers. Indeed, VASP recruitment to FAs, although limited, can occur independently of zyxin (Hoffman et al., 2006). Zyxin contributes to adhesion maturation and tension regulation, localizes to FAs and stress fibers via its LIM domains (Hirata et al., 2008) and is crucial for targeting VASP to these sites (Hansen and Beckerle, 2006; Hoffman et al., 2006; Wang et al., 2019). Our cryo-ET results show that zyxin deletion exacerbates the VASP^-/-^ phenotype, displacing VASP from adhesion sites and compromising the tension applied at FAs, as zyxin is key to actomyosin stress fiber maintenance and repair (Smith et al., 2010). This likely reflects disengagement of the ‘molecular clutch’ and FA disassembly due to insufficient contractile force (Hoffman et al., 2006; Smith et al., 2010). Consistently, double knockout cells lacking both VASP and zyxin displayed additive alterations, including more severe disruption of adhesion morphology and actin organization. Altogether, these findings indicate that VASP is essential for supplying aligned actin filaments to forming adhesions near the plasma membrane, an early step in FA maturation and stress fiber formation, while zyxin, via tension-mediated feedback, reinforces filament density and organization, thereby supporting further adhesion maturation and dynamics.

### A confined dorsal tropomyosin association with stress fibers defines a pool of stable actin filaments for contractile integration

A key feature of the actin cytoskeleton is its ability to assemble into functionally distinct structures, often associated with specific, exclusive sets of binding proteins (Pollard and Cooper, 2009). Tropomyosins, a diverse and conserved family of actin-binding proteins (Geeves et al., 2015), is recruited to lamellipodial actin and adhesions (Gunning et al., 2015). Tropomyosin isoforms such as 1.6 and 3.1 have been shown to localize to FAs and may segregate to distinct FA subdomains (Kumari et al., 2024). Tropomyosin isoforms were also shown to modulate the activity of actin-binding proteins at adhesion sites. Isoforms1 and 2 can protect actin filaments against cofilin-mediated severing, whereas isoform 4 promotes the coupling of adhesion-associated F-actin to contractile dorsal stress fibers. This facilitates adhesion strengthening through myosin-generated tension while simultaneously suppressing vectorial actin assembly via VASP inactivation through phosphorylation (Gateva et al., 2017; Tojkander et al., 2015; Tojkander et al., 2011). Our cryo-ET analysis revealed that tropomyosin decoration is enriched in the dorsal portion of the FA-associated actin network and remains unaffected by changes in actin organization or tension induced by the individual and combined removal of VASP and zyxin. These findings identify a distinct subpopulation of tropomyosin-decorated actin filaments, likely representing a more stable filament pool that may preferentially interact with myosin and facilitate engagement of FA-associated actin with the contractile actomyosin machinery.

In summary, our study suggests that VASP and zyxin play distinct yet complementary roles in organizing focal adhesion-associated actin architecture. These findings indicate that actin filament specialization within adhesions is achieved through the coordinated yet independent regulation of filament orientation and density, and tropomyosin association, offering new insight into how cytoskeletal architecture is tuned to support adhesion maturation and mechanotransduction. With continued advances in cryo-tomogram image processing, it is anticipated that the spatial distribution of additional actin binding partners at FAs will be resolved, ultimately enabling the reconstruction of a high-resolution 3D map of the adhesion machinery.

### Materials and methods Generation of knockout cell lines

The VASP^-/-^, zyx^-/-^, and DKO MEFs were generated from the parental MEF cell line expressing vinculin-Venus (Grashoff et al., 2010) following published protocols (Ran et al., 2013; Weber et al., 2021). The gRNAs were designed and selected on Benchling (https://www.benchling.com/) with the target sequences: 5’-CTCGGAAGGAGTTAGCAGTG – 3’ and 5’ – CGGGGCGAACTTCTTCTGCG – 3’ on exon 2 of VASP and zyx. The gRNAs were constructed into pSpCas9(BB)−2A-GFP (pX458) and transfected into MEFs by jetOPTIMUS^®^ DNA transfection reagent following the manufacturer’s protocol. The GFP-positive cells were sorted 24 hr after transfection for clone culture. To verify mutate clones, the genomic DNA of the candidate clones was extracted with the GenElute Mammalian Genomic DNA kit (Sigma-Aldrich). The fragments containing the expected indel mutation were amplified by PCR and sent for Sanger sequencing. Next, the sequencing results were analyzed with TIDE(Brinkman et al., 2014) to identify the KO clones. The fragments were further cloned and sequenced to verify the mutation (Fig. S1a). The zyxin-/- was then used to generate the DKO with the same CRISPR construct targeting VASP. Eventually, the expression of VASP and zyxin of the selected knockout cell lines were analyzed by western blotting and immunofluorescence (Fig. S1b and S1c).

### Cell culture and live-cell fluorescence microscopy

Cells were seeded on glass bottom plates covered with 10ug/ml fibronectin and were maintained at 37°C in DMEM supplemented with 10% FCS under a humidified atmosphere of 5% CO2. Time-lapse movies were recorded using the Real Time Delta Vision System (Applied Precision, LLC), which consists of an inverted microscope (IX71; Olympus) equipped with a CoolSnap HQ camera (Photometrics) and weather station temperature controller (Precision Control, LLC), operated by SoftWoRx and Resolve3D software (Applied Precision, LLC). Images were acquired with a Plan-Apochromat 60×/1.40 NA objective (Olympus). Image stacks were acquired at 1-minute intervals.

Adhesion pixels from raw movies were classified using Ilastik software (Berg et al., 2019), exported to Fiji/ImageJ2 software as segmented pixels and FA particles were further segmented using the Trainable Weka Segmentation plugin. FA number, size, and fluorescence intensity, along with the overall cell area, were quantified. FA fluorescence intensity was measured in MEFs expressing vinculin-Venus at levels comparable to endogenous vinculin, which had been deleted (Grashoff et al., 2010). To enable quantitative comparison, all images were acquired using identical illumination settings. Data analysis and figure preparation were carried out using MATLAB R2019b and GraphPad Prism 10.

Temporal ratio images were calculated and presented as described previously (Zamir et al., 2000). In this process, the intensity ratio of identical pixels in images acquired at different time points is calculated and represented by a color scale, such that “new pixels” are blue, pixels that disappeared “lost pixels” are red, and persistent pixels are green.

Autocorrelation analysis was performed as described previously (Zaidel-Bar et al., 2007b), comparing the relative intensities of all the image pixels at two time points. Temporal ratio images and autocorrelation analysis were performed on segmented images using Ilastik (Berg et al., 2019) for optimal visualization of FA. The autocorrelation values are the average of 6 independent movies for each phenotype. The exponential decay time +/- standard deviation was calculated, as well as the P-values from the Beta function for the Student T-test.

### Micropatterning essay and light microscopy

We used the adapted methods of micropatterning essay for cells (Boujemaa-Paterski et al., 2020). Coverslips (CVs) were used to assemble the reaction chambers were drastically cleaned by successive chemical treatments: 2 hr in 2% Hellmanex III, rising in ultrapure water, 30 min in acetone, 30 min ethanol 96%, rinsing in ultrapure water. Slides and CVs were dried using a filtered nitrogen gas flow and oxidized with oxygen plasma (3 min, 30 Watt, Femto low-pressure plasma system Type A, Diener electronic GmbH, Germany) just before overnight incubation in a solution containing tri-ethoxy-silane-PEG (5 kDa, PLS-2011, Creative PEGWorks, USA) 1 mg/mL in ethanol 96% and 0.02% of HCl, with gentle stirring. mPEG-silane-coated CVs were then rinsed in ethanol and extensively in ultrapure water. Passivated CVs were then stored in a clean container and used within a week time.

Cells were confined to predefined locations on glass coverslips using a micropatterning strategy that consisted of printing adhesive patterns onto a protein-repellent surface. PEG-coated CVs were exposed to short-wavelength UV radiation (184.9 nm and 253.7 nm, Jelight, USA) for 2 min through transparent micropatterns printed on a photomask (Compugraphics, Germany). Immediately after UV exposure, micropatterned CVs were incubated in 10 μg/ml fibronectin solution for 15 min at room temperature under gentle agitation, then washed in PBS buffer. CVs were mounted in a Ludin chamber (Elveflow, France) prior to cell seeding. Cells were allowed to adhere on the micropatterns for 3 hours, then washed in 37°C-warmed phosphate saline solution (PBS) before being fixed in 3.7% paraformaldehyde (PFA). After removal of the fixative, nuclei were stained with 5 µg/mL Hoechst 33342 dye (B2261, Sigma) for chromatin and the actin cytoskeleton with one unit of Alexa Fluor® 647 phalloidin (A12387, ThermoFisher Sc.). Labelled fixed cells were then embedded in ProLong Glass antifade (Thermofisher). For epifluorescence imaging, we used the inverted, widefield Leica Thunder imager that consists of DMi8 microscope equipped two sCMOS B/W camera (Leica monochrome fluorescence DFC9000 GTC) operated by LAS X software. Images were acquired with a Highly Corrected Plan Apochromat, HC PL APO PH3 100×/1.4 NA objective (Leica).

Focal adhesion (FA) particles and entire micropatterned cells were independently segmented using the Trainable Weka Segmentation plugin (Fiji/ImageJ2). FA number, size, and fluorescence intensity, along with the overall cell area, were quantified. FA fluorescence intensity was measured in MEFs expressing vinculin-Venus at levels comparable to endogenous vinculin, which had been deleted (Grashoff et al., 2010). To enable quantitative comparison, of all images were acquired using identical illumination settings. Data analysis and figure preparation were carried out using MATLAB R2019b and GraphPad Prism 10. To calculate average fluorescence intensity of FAs in individual cells, we separately piled images from the vinculin-Venus and Alexa647-labelled actin cytoskeleton channels. Piled images were then aligned according to the circular geometry of patterned cells using ‘Template Matching’ plugins of Fiji software. Average projection images, representing the pixel-wise mean intensity across the stack, were generated using Fiji’s default functions.

### Immunofluorescence

MEF cells were fixed with 3.7% paraformaldehyde. After washing and permeabilization with 0.1% Triton-X/PBS (3 x 5 min), cells were blocked with 1% BSA in PBST (1x PBS, 0.1% Tween-20) for 1 hr. Primary antibodies were incubated for 1 h, followed by extensive washing with PBST and incubation with respective secondary antibodies for 1 h. For immunolabeling of Arp 2/3 complex, cells were incubated with rabbit polyclonal anti-p34-Arc (1:100, Sigma–Aldrich 07-227), then with goat-anti-rabbit IgG H&L Alexa Fluor 488 (1:400, Abcam ab150077) and Alexa Fluor 647 phalloidin (1:100, Thermo Fisher Scientific. A22287). Prior to imaging, coverslips were mounted with fluorescence mounting medium (Dako Omnis). Confocal images were acquired with Olympus IXplore SpinSR10 super-resolution imaging system equipped with two sCMOS cameras and processed with ImageJ (Schneider et al., 2012). For immunolabeling of VASP and zyxin, cells were incubated with rabbit polyclonal anti-VASP (1:1000, gift of Jan Faix, (Damiano-Guercio et al., 2020)) and mouse monoclonal anti-zyxin clone 164D4 (1:1, gift of Klemens Rottner, (Rottner et al., 2001)). These primary antibodies were detected with polyclonal secondary Alexa Fluor 647 donkey anti-rabbit (1:500, Thermo Fisher Scientific A31573) and Alexa Fluor 647 donkey anti-mouse (1:500, Jackson ImmunoResearch 715-605-150) antibodies. The cells nuclei were stained by Hoechst reagent (1:100, Sigma Aldrich B2261). Imaging was performed with an inverted widefield IX83 Olympus microscope equipped with an Olympus planApo N 60x/1.43 NA oil objective and a Hamamatsu ORCA-FUSION digital camera. The system was operated by Cellsens software (Olympus).

### Western blotting

For preparation of total cell lysates, MEFs were cultured to ∼70% confluency, washed once with warm PBS, and trypsinized. Cell pellets were washed twice with cold PBS and lysed with cold lysis buffer (1x PBS, 1% Triton X-100, cOmplete EDTA free protease inhibitor cocktail (Roche Diagnostics 52434800)) for 1 h on ice. Cell lysates were subsequently homogenized by passing them 10 times through a 27G syringe (Eclipse Needle, 302436 BD) and incubated 1 h on ice. Protein concentrations were determined by Bradford assay (Bio-Rad 500-0006) using an Ultrospec 10 Cell Density Meter (Biochrom) at 595 nm. Equal amounts of protein (30 µg per lane) were subjected to SDS–PAGE and transferred by wet blotting onto PVDF membranes (Immobilon-P, Millipore IPVH00010). Membranes were blocked with PBST (1X PBS, 0.05% Tween-20) containing 5% Nonfat dried milk powder (AppliChem A0830) for 1 h, then incubated with primary antibodies for 1 h in the same buffer. After extensive washing of membranes in PBST and incubation with secondary peroxidase-conjugated antibodies for 1 h, blots were developed with Immobilon Western Chemiluminescent HRP Substrate (Millipore P90720), according to manufacturer’s instructions, using a Fusion-FX7 (Vilber). Primary antibodies were rabbit polyclonal anti-VASP (1:100, gift of Jan Faix, (Damiano-Guercio et al., 2020)) and mouse monoclonal anti-zyxin clone 164D4 (1:1, gift of Klemens Rottner, (Rottner et al., 2001)), anti-α-tubulin (1:200,000, Sigma-Aldrich T5168). The polyclonal secondary peroxidase-conjugated antibodies were donkey anti-mouse HRP (1:5000, Jackson Immuno Research 715-035-150) and donkey anti-rabbit HRP (1:5000, Jackson Immuno Research 711-035-152).

### Cell culture and correlative light electron microscopy

WT and knockout MEFs expressing vinculin-venus (Grashoff et al., 2010; Martins et al., 2021; Ringer et al., 2017) were used to show FA and optimal adhesion. The cells were cultured in Dulbecco’s Modified Eagle’s Medium (Sigma–Aldrich, D5671) with 10% fetal bovine serum (Sigma–Aldrich, G7524), 2 mM L-glutamine (Sigma–Aldrich, G7513), and 100 μg/ml penicillin-streptomycin (Sigma– Aldrich, P0781) at 37 °C and 5% CO_2_. Glow-discharged EM grids with carbon support film (R2/2, Au mesh, Quantifoil, Jena, Germany) were coated with 10 μg/ml fibronectin, 2 hr room temperature. The cells were detached with 5 mM EDTA, resuspended in serum-free medium, and seeded onto the EM grids. The cells were then incubated at 37 °C and 5% CO_2_ for 3 hr to allow FA formation. The grids were then transferred into 1x PBS (Fisher Scientific, BP399-1) in a 35×14 mm glass-bottom cell culture dish (MatTek, P35G-0-14-C). For CLEM, a low magnification overview and 16 high magnification images around the center of each grid were taken with an automated inverted microscope (DMI4000 B, Leica Microsystems, Wetzlar, Germany) equipped with a fluorescence lamp and a monochromatic digital camera (DFC365 FX, Leica Microsystems), and operated with Leica LAS X software (Leica Microsystems, GmbH, Germany). Before plunge-freezing in liquid ethane, 4 ml of BSA-coated 10 nm fiducial gold markers (Aurion, Wageningen, Netherlands) was applied onto the grids. The entire procedure was carried out within 10 min to ensure the optimal preservation of each sample.

### Cryo-electron tomography

A Titan Krios G2 transmission electron microscope (Thermo Fisher Scientific, Waltham, MA) equipped with an energy filter and a K2-summit direct electron detector (Gatan, Pleasanton, CA) was used for data acquisition. The microscope was operated at 300 keV in zero-loss mode; the energy filter slit width was set to 20 eV. The microscope was controlled by SerialEM(Mastronarde, 2005). Tilt series were acquired at bidirectional or dose symmetric schemes, ranging from −60 to 60° with 3° increments at −4 μm defocus. The magnification was set to ×64,000 resulting in a pixel size of 2.21 Å. The dose-fractionated mode was used, and the accumulated electron dose per tilt series is around 130 e^-^/Å^2^. The tomograms were reconstructed and CTF-corrected with IMOD(Kremer et al., 1996).

### Actin and tropomyosin-actin filament 3D reconstruction and polarity determination

We followed the APT framework for filament 3D structure reconstruction and polarity determination (Chung et al., 2022). First, we extracted 2,341,836 actin particles from 186 tomograms with CrYOLO (Wagner et al., 2019). The inter-particle distance was set to 5.3 nm along the filament for optimal 3D filament tracing. We then exported the filament coordinates to IMOD for subtomogram reconstruction. The central 11 nm of each subtomogram was projected before 2D classification and 3D refinement with RELION (Scheres, 2012). Eventually, 661,369 actin filament segments were used for filament polarity determination.

To reconstruct the tropomyosin-actin filament, an iterative particle selection and classification process was employed. Initially, particles displaying tropomyosin features were identified as tracers to pinpoint adjacent particles on the same filaments. Then, the enriched particle pool was subjected to additional 2D classification and subset selection. This iterative process was repeated until no further improvements in 2D classification were observed. Subsequently, the selected particles were subjected to 3D refinement followed by 3D classification without alignment. Ultimately, 37,330 particles were utilized for the final refinement of the tropomyosin-actin filament reconstruction.

To determine the polarity of the actin filaments, an adapted version of APT was used to determine the filament orientation corresponding to the FA. In each tomogram, filaments identified with CrYOLO were exported and defined as *F*_j_^i^ with *i* the filament index and *j* the segment index. On each filament, the 3D coordinates (*C*_j_^i^) of the segments were then used for a polynomial regression fit for the filament model (*f*(*C*_j_^i^)). The tangent vector of each segment (*T*_j_^i^) derived from the model were then used as the observed polarity. We then compare the polarity of each segment (*P*_j_^i^) to their tangent vectors. The segments following their tangent vectors within 30 degrees were given a direction labeled 1 and else 0 (ε*F*_j_^i^);

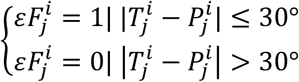

The *mean*(ε*F*_j_^i^) were used as the combined confidence score for screening the proper polarity assignment (*ccs*(*F*^i^) ≥ 0.6) (Chung et al., 2022; Martins et al., 2021).

In each tomogram *t*, the FA is defined as an ellipse and its orientation is defined by the major axis *A*^t^. The orientation toward the cell margin is defined as 0 degree. The relative orientation of each actin segment to the FA (*O*_j_^i^) was then calculated by comparing its polarity of the major axis in the x-y plane:

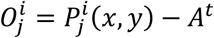

The overall filament to FA orientation is calculated by the average of all the segment-FA orientations (*mean*(*O*_j_^i^)). For general categorization of filament-FA orientation, we defined the 3 intervals:

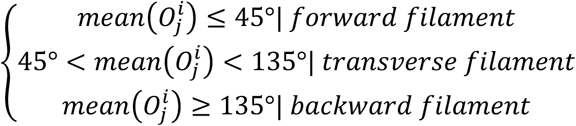

### Tomogram visualization

All isosurface visualizations of actin filaments were rendered using UCSF ChimeraX (Pettersen et al., 2021). The refined structures of actin filament and tropomyosin-actin filament were used for visualizing the orientation of each respective segment analyzed by APT. Segments on the same filaments share the same color regarding their averaged angles. Some parts of the filaments were given transparent colors, where the segments were locally poorly resolved and excluded from the APT analysis, while the filament polarity can still be confidently assigned.

## Quantification and statistical analysis

The modified plotting toolbox (Morel, 2018) was used for all the boxplots and violin-plots. The significant difference in data was tested by one-way ANOVA and the *multcompare* function using the Tukey-Kramer method in MATLAB. To compare the significant difference of filament distribution in violin-plots of each two groups, a 10000 times permutation test was designed to calculate the 95% confidence intervals of the 5 and 95 data percentiles as the edges of the distribution. Only *P*<0.05 in both edges is considered as a significantly different distribution.

## Statistics and reproducibility

We acquired 186 cryo-tomograms on 45 WT cells, 8 VASP KO cells, 29 zyxin KO cells, and 13 DKO cells. For actin filament analysis, 57 tomograms of WT-distal region, 9 of WT-proximal region, 19 of VASP KO cells, 44 of zyxin KO cells, and 21 of DKO cells containing the full set of image and low residual errors were used.

## Data availability

The EM structures of actin filament and tropomyosin-actin filament were uploaded on the Electron Microscopy Data Bank with the accession codes EMD-17742 and EMD-17744, respectively. A tomogram of WT, zyxin^-/-^, VASP^-/-^, and DKO were uploaded on the Electron Microscopy Data Bank with the accession codes EMD-17745, EMD-17746, EMD-17750, and EMD-17752, respectively. The ATP code is available at https://github.com/WChung2/actin_polarity_toolbox_adhesion.

## Contributions

W.L.C. prepared and acquired the tomograms and analyzed the data with the help of M.E. R.B.P. contributed to patterned cell assay and analysis. O.M. conceived the work and financed the project. R.B.P wrote the manuscript with contributions from all authors.

## Acknowledgments

This work was funded by the Swiss National Science Foundation (SNSF 310030_207453) to O.M. and by the European Research Council (810057-HighResCells). We thank Prof. Jan Faix (Hannover Medical School, Germany) and Prof. Klemens Rottener (Helmholtz Centre for Infection Research, Braunschweig, Germany) for their help with VASP and zyxin antibodies. We also thank the Center for Microscopy and Image Analysis (ZMB) at the University of Zurich, as well as Sara Billeter and Ginevra Destefani for assistance with western blotting and immunofluorescence imaging.

