## Supplementary figures for "VASP regulates the polar organization of adhesion-associated actin filaments"

**Keywords**: actin, VASP, zyxin, cytoskeletal organization, focal adhesion, cryo-ET

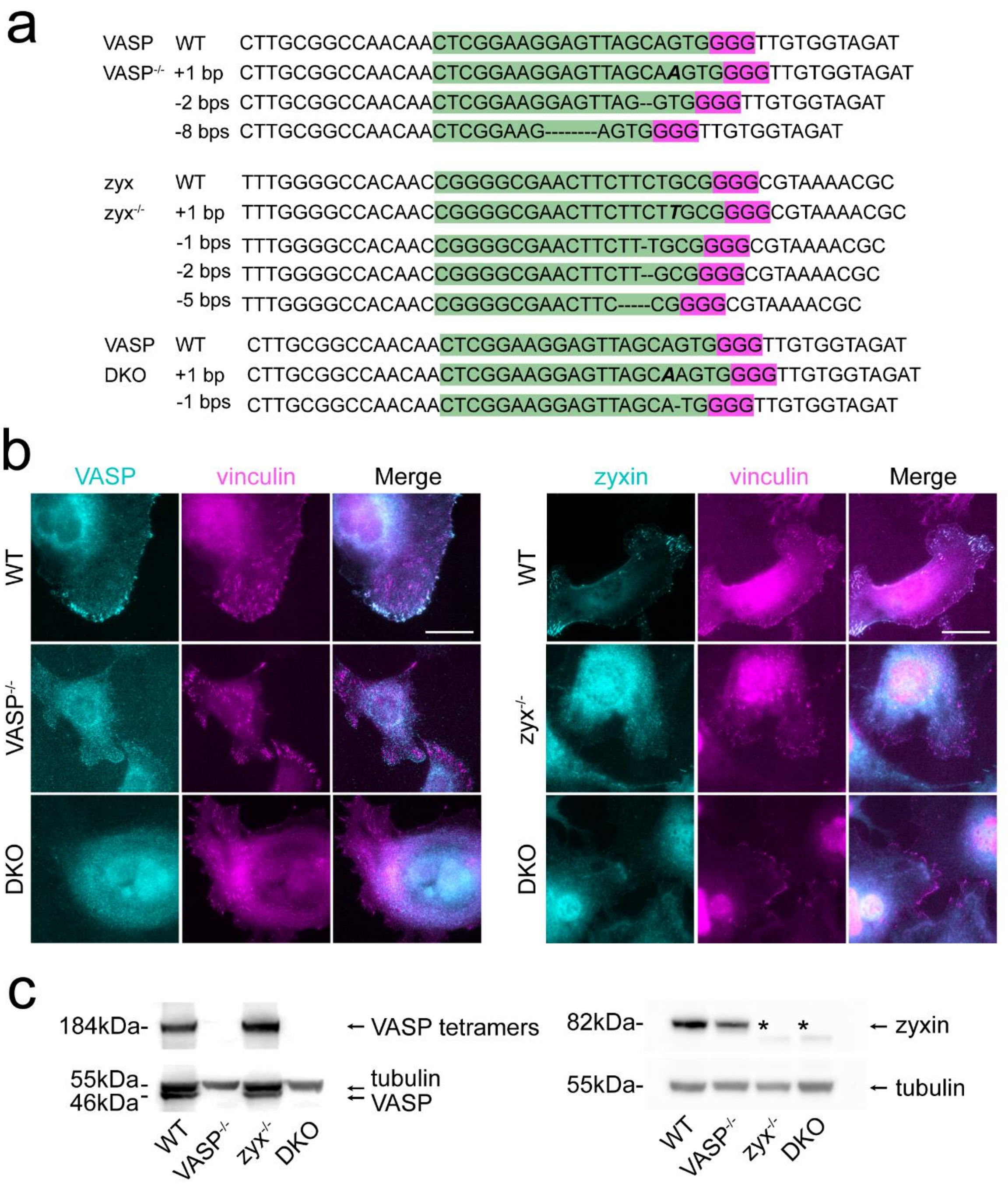


Figure S1. **Genotyping and validation of VASP$^{-/-}$, zyxin$^{-/-}$, and double knockout (DKO) MEFs. a,** Genomic sequences of WT and KO cells at the CRISPR-Cas9 target sites. Guide RNA binding regions and Protospacer Adjacent Motif (PAM) sequences following the target site are highlighted in green and magenta, respectively. **b,** Representative immunofluorescence images showing VASP and zyxin expression in WT and KO cells. Scale bar, 10 μm. **c,** Representative immunoblots of VASP and zyxin expression in WT and KO cells. α-tubulin serves as a loading control. * : zyxin fragments are detectable due to truncated expression resulting from a nonsense mutation downstream of the N-terminal knockout sites. (related to Figure 1)

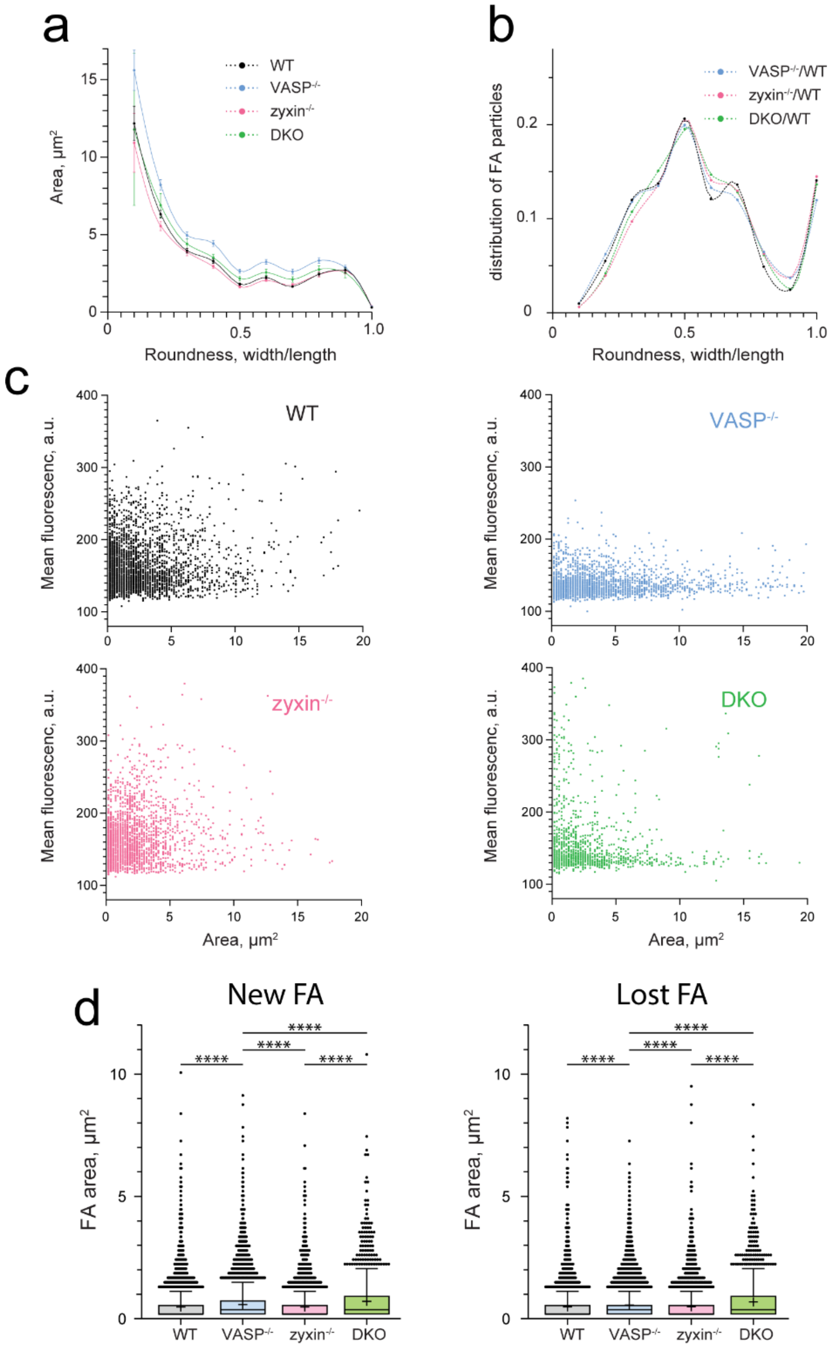


Figure S2. **Quantitative analysis of adhesion particles in wildtype and KO cells. a-c,** Adhesion particles were segmented using Ilastik (Berg et al., 2019) and built-in plugins in Fiji (ImageJ), and analyzed in MATLAB. No significant differences were observed between cell lines in the variation of adhesion area as a function of roundness (a), the distribution of adhesion roundness (b), or vinculin mean fluorescence relative to adhesion area (c). **d,** Comparison of the distribution of newly formed or lost adhesion particles across WT and KO cell lines revealed statistically significant differences, although changes in median (cross) and mean (bar) values were negligible. Statistical analysis was performed using one-way ANOVA and Holm–Šídák multiple comparisons test; p-values are indicated as **** < 0.0001. (Related to Figure 1)

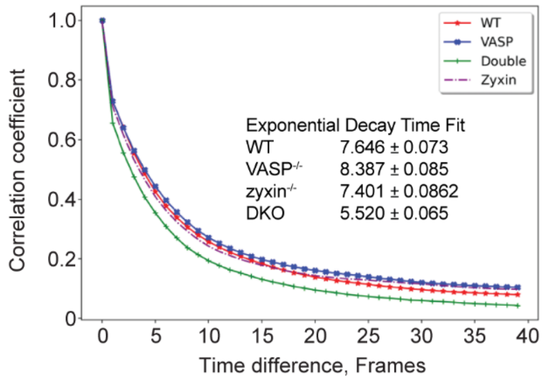


**P-values from Beta function for the Student T-test**

| | WT | VASP-/- | zyxin-/- | DKO |
|---|---|---|---|---|
| WT | | 9.244 E-06 | 1.236 E-01 | 1.388 E-25 |
| | VASP-/- | | 1.101 E-07 | 2.710 E-31 |
| | | zyxin-/- | | 3.150 E-20 |
| | | | DKO | |

Figure S3. **Dynamics of adhesion-associated vinculin assessed by autocorrelation analysis.** Relative intensities of all adhesions pixels at two time points were compared. Autocorrelation analysis was performed on segmented images using Ilastik (Berg et al., 2019). Values represent the average of six independent movies per phenotype. Exponential decay times are reported ± standard deviation, and statistical significance was determined using the Beta function for the Student's t-test (Zaidel-Bar et al., 2007b; related to Figure 1).

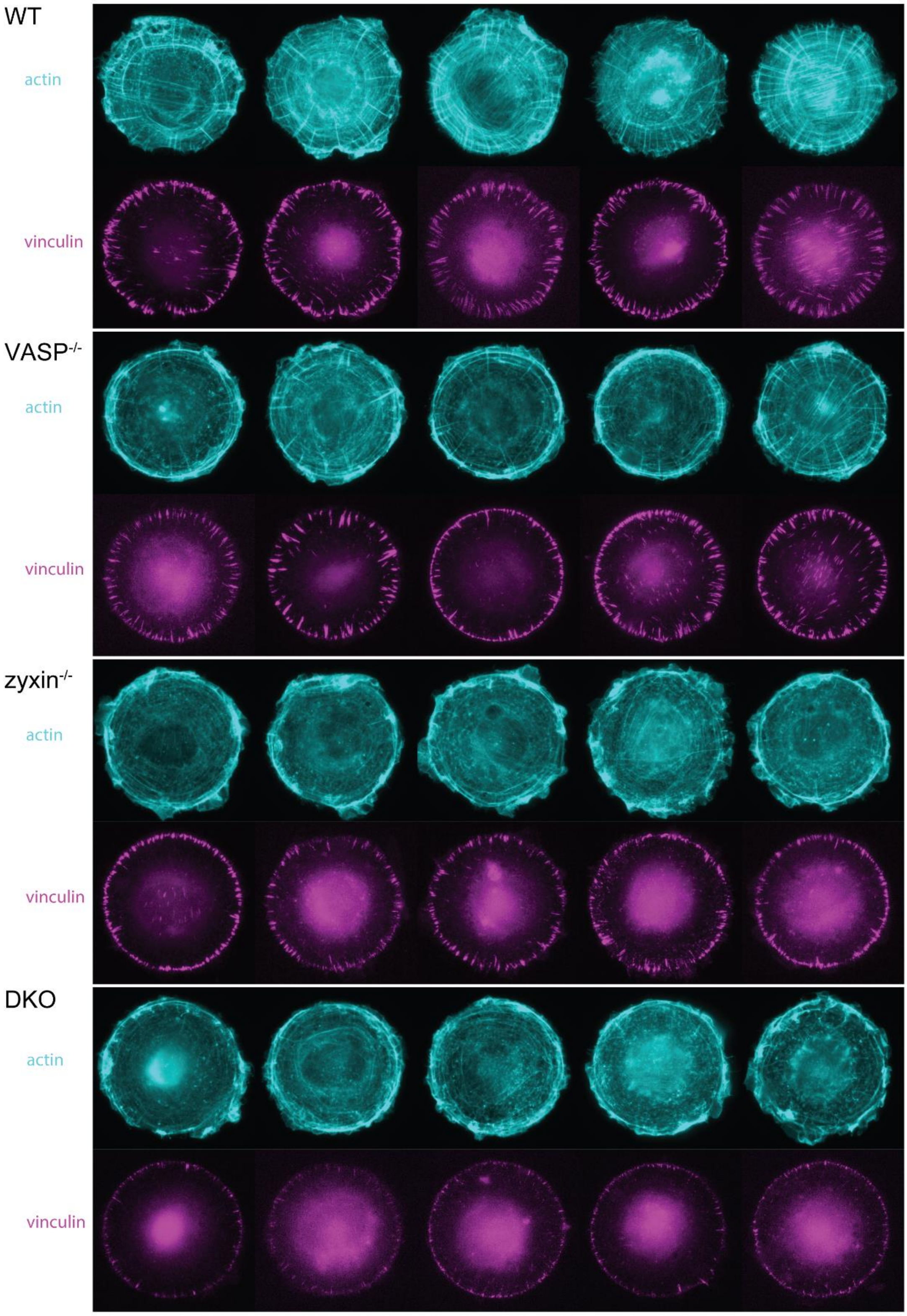


Figure S4. **Loss of VASP and zyxin affects FA, actin network organization, and membrane protrusions of cells on patterned surfaces.** Five representative images of MEFs spread on circular micropatterns (48 µm in diameter, ~1810 µm$^2$), selected from a total of 112 for WT, 104 for VASP$^{-/-}$, 129 for zyxin$^{-/-}$, 96 for DKO cells analyzed. Vinculin and actin are shown in magenta and cyan, respectively. (related to Figure 2).

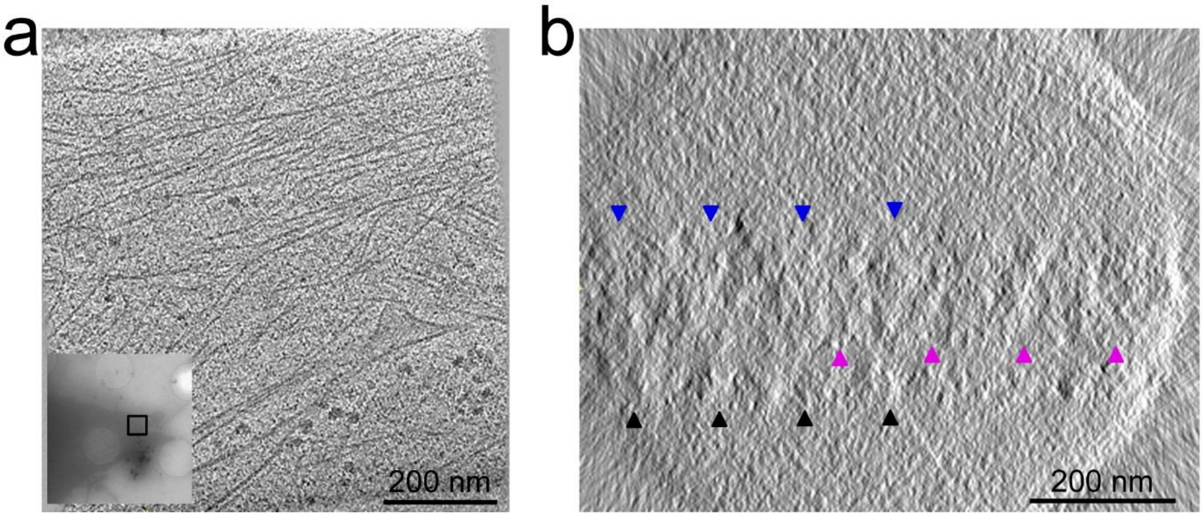


Figure S5. **Determination of substrates within tomograms. a,** Representative x–y slice of a tomogram at a focal adhesion. A corresponding low-magnification correlative image is shown at the bottom left. **b,** x–z slice of the tomogram shown in a. The substrate beneath the cell on the EM grid is marked by magenta arrowheads. Blue and black arrowheads indicate the top of the sample and the opposite side of the EM grid, respectively. (related to Figure 3)

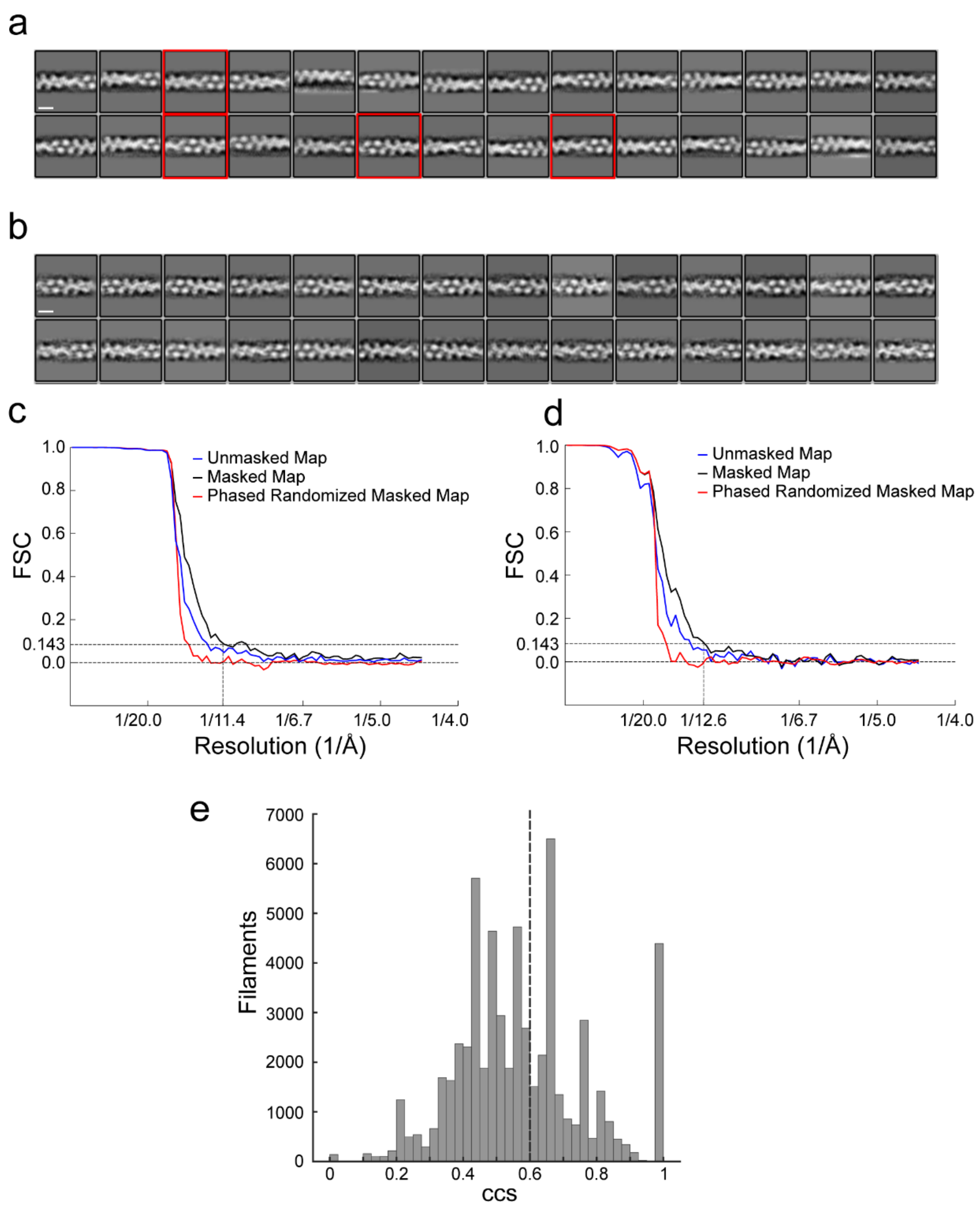


Figure S6. **Reconstruction of actin and tropomyosin–actin filaments by Actin Polarity Toolbox (APT). a,** Representative two-dimensional (2D) class averages of actin segments. Classes displaying tropomyosin features are highlighted in red and used for further reconstruction of tropomyosin–actin. Box size, 36 nm; scale bar, 8 nm. **b,** Representative two-dimensional class averages of tropomyosin–actin segments. Box size, 36 nm; scale bar, 8 nm. **c,** Refined actin structure (corresponding to Fig. 3a) with a spatial frequency of 1/11.4 Å, determined using the 0.143 gold-standard Fourier shell correlation criterion. **d,** Refined tropomyosin–actin structure (corresponding to Fig. 3d) with a spatial frequency of 1/12.6 Å, determined using the 0.143 gold-standard Fourier shell correlation criterion. **e,** Combined confidence score of the acquired dataset, as described in Methods. Approximately 42% of filaments passed the 0.6 confidence score threshold (N = 186). (related to Figure 3)

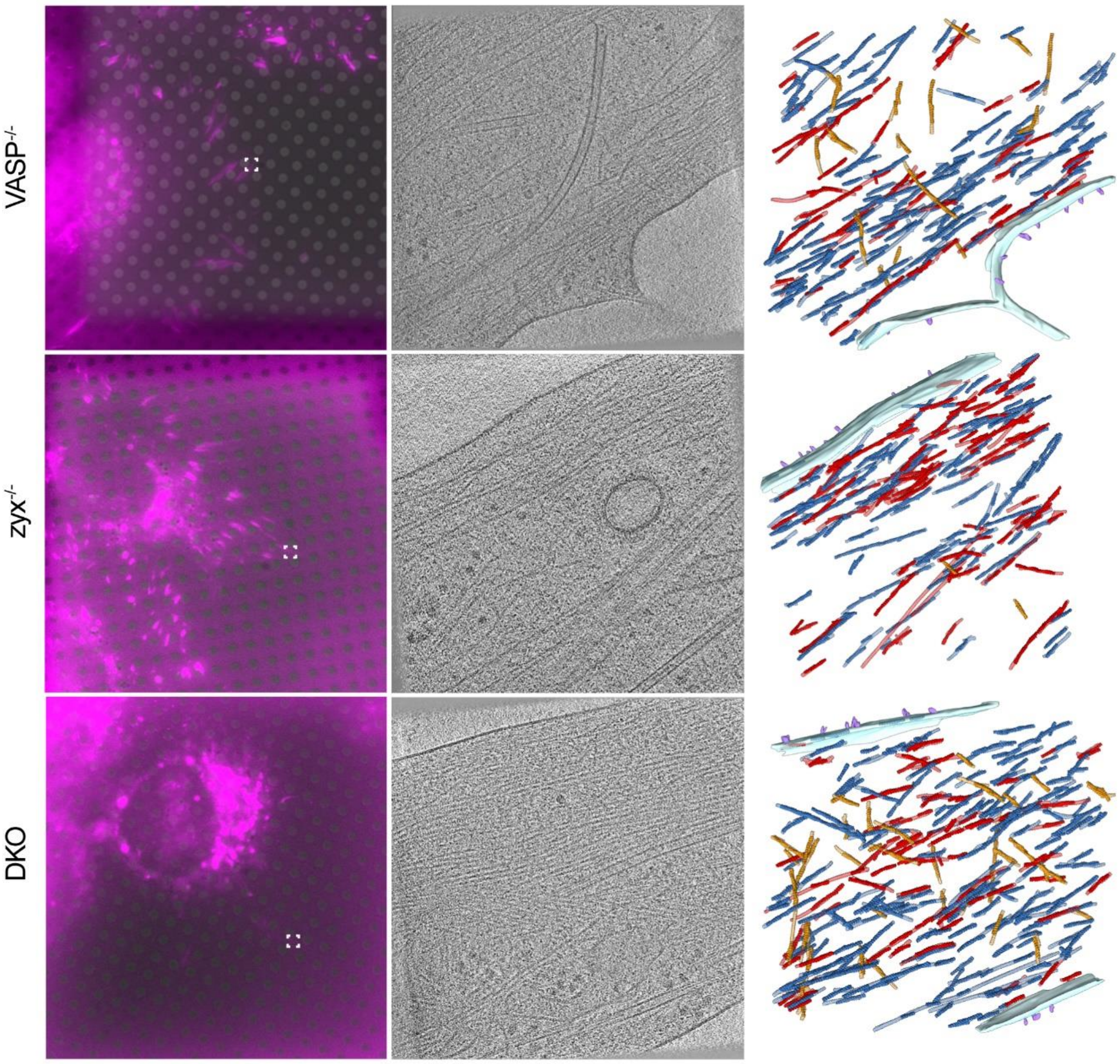


Figure S7. **Correlative light and electron microscopy (CLEM) of knockout (KO) cells.** Representative fluorescence images of KO cells expressing vinculin on electron microscopy (EM) grids are shown in the left panel. The middle panel presents an x–y slice of each tomogram acquired at the indicated regions (white boxes). The right panel displays isosurface renderings of the corresponding tomograms. Filaments are colored as in Fig. 4c: where forward filaments (blue) have their barbed ends oriented toward the cell periphery along the FA axis; backward filaments (red) are oriented in the opposite direction; transverse filaments (brown) are not aligned with the FA axis. The plasma membrane was manually segmented and is shown in light blue. (related to Figure 5)

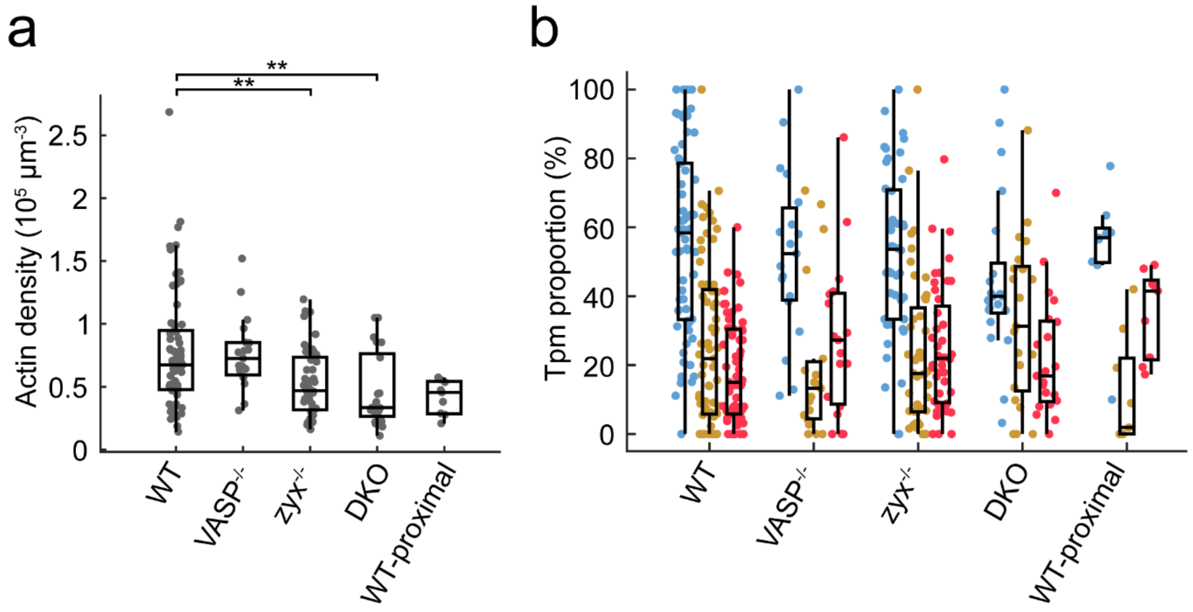


Figure S8. **Actin filament density and tropomyosin distribution at focal adhesions in wild-type (WT) and knockout (KO) cells. a,** Actin filament density at FAs in WT and KO cells. Parameters were measured in all cells, 54 tomograms for WT, 19 for VASP$^{-/-}$, 44 for zyxin$^{-/-}$, 21 for DKO. Statistical comparisons using the Tukey-Kramer method, and a one-way analysis of variance (ANOVA) showed significant variations among the measured parameters, **P < 0.01. **b,** Proportion of tropomyosin associated with forward (blue), parallel (brown), and backward (red) filaments at FAs in WT and KO cells. WT-proximal represents the distributions observed in the proximal region of the adhesion particle defined by CLEM, as described in Fig. 5. (related to Figure 5)
